# Amygdala Subregion Volumes and Apportionment in Preadolescents — Associations with Age, Sex, and Body Mass Index

**DOI:** 10.1101/2024.10.07.617048

**Authors:** L. Nate Overholtzer, Carinna Torgerson, Jessica Morrel, Hedyeh Ahmadi, J. Michael Tyszka, Megan M. Herting

## Abstract

The amygdala, a key limbic structure, is critical to emotional, social, and appetitive behaviors that develop throughout adolescence. Composed of a heterogeneous group of nuclei, questions remain about potential differences in the maturation of its subregions during development. In 3,953 9- and 10-year-olds from the Adolescent Brain Cognitive DevelopmentlZI Study, the CIT168 Amygdala Atlas was used to segment nine amygdala subregions. Linear mixed-effects models were used to examine the effects of age, sex, pubertal stage, and body mass index z-score (BMIz) on subregion volumes and their relative apportionment within the amygdala. Distinct associations were observed between age, sex, and BMIz and whole amygdala volume, subregion volumes, and subregion apportionment. Pubertal stage was not related to amygdala subregion volumes. Age was associated with near-global expansion of amygdala subregions during this developmental period. Female sex was linked to smaller volumes in most amygdala subregions, with larger relative apportionment in the dorsal subregions and smaller apportionment in the basolateral ventral paralaminar subregion. Higher BMIz was associated with smaller volumes in large basolateral subregions, with increased relative apportionment in smaller subregions. These findings provide a foundational context for understanding how developmental variables influence amygdala structure, with implications for understanding future risk for brain disorders.

**Highlights:**

- Segmentation of amygdala subregions in nearly 4,000 preadolescents.
- Age, but not puberty, was associated with a near-global expansion of the amygdala.
- Sex differences exist in preadolescent amygdala apportionment.
- Childhood obesity is linked to differences in the basolateral amygdala.

## Introduction

Adolescence marks a critical period of substantial developmental changes to the amygdala (Dennison et al., 2013; Scherf et al., 2013), coinciding with the maturation of complex social and emotional behaviors that develop between childhood and adulthood (Blakemore, 2008). Although neuroimaging studies typically consider the human amygdala as a homogenous subcortical region, it can be hierarchically organized into three major subdivisions — basolateral, centromedial, and superficial (or “cortical-like”) nuclear groups (Kedo et al., 2017; LeDoux, 2007; Tyszka and Pauli, 2016) — and further decomposed into thirteen individual nuclei. These nuclei exhibit distinct cytoarchitecture, connectivity, and functional roles (Amunts et al., 2005; Kedo et al., 2017; McDonald, 1998), while sharing similarities in cytoarchitectonic features to other nuclei of the same group.

The basolateral nuclear group (BLN) — containing the lateral (LA), basolateral (BL), basomedial (BM), and paralaminar (PL) nuclei — is the primary afferent layer for processing high-level sensory input in the amygdala and primarily projects to the central (CEN) nucleus (Hintiryan et al., 2021; LeDoux, 2007). The BLN, through its connections with the prefrontal cortex, is important for emotional learning and memory (McDonald, 2020). The paralaminar nucleus (PL) is unique in that it also exhibits characteristics similar to cortical-like groups (Kedo et al., 2017), and it contains late-maturing neurons that migrate during the pubertal period (Page et al., 2022). The central nucleus (CEN), cortical and medial nuclei (CMN), and anterior amygdaloid area (AAA) constitute the centromedial nuclear group and integrate this sensory information to mediate behavioral and autonomic responses (Bzdok et al., 2013). The CEN is the recipient of most intrinsic amygdala projections and, accordingly, the primary output nuclei of the amygdala in a striatum-like system, where it plays a critical role in mediating defensive and appetitive behaviors (Fadok et al., 2018; LeDoux, 2007). The CEN also directly projects to the paraventricular nucleus of the hypothalamus, where it can modulate cortisol release as an extended component of the hypothalamic-pituitary-adrenal axis (Gray et al., 1989). Finally, the superficial nuclear group includes the amygdala transition area (ATA), which contains both the entorhinal cortex and hippocampal boundaries, and the amygdalostriatal transition area (ASTA) (Kedo et al., 2017; Tyszka and Pauli, 2016). The functional roles of the superficial nuclear group remain poorly understood; however, functional MRI has linked the region to processing social relevance and emotional valence (Goossens et al., 2009; Koelsch et al., 2013).

Structural magnetic resonance imaging (sMRI) studies have shown that total amygdala volumes follow a curvilinear growth trajectory, beginning in late childhood and continuing through adolescence (Dima et al., 2022; Herting et al., 2018). Sex differences in total amygdala volumes become more substantial throughout adolescence, with females displaying smaller volumes that peak relatively earlier (∼14 years) than males who undergo a more prolonged growth period (∼18-24 years) (Dima et al., 2022; Herting et al., 2018). However, age poorly explains variability in amygdala volume compared to other subcortical regions (Dima et al., 2022), suggesting substantial inter-individual variability in amygdala development. Other developmental factors, such as pubertal progression (Scherf et al., 2013) and indicators of obesity (Kim et al., 2020; Perlaki et al., 2018), may also contribute to changes in total amygdala volumes during childhood and adolescence. Beyond the whole amygdala, assessing developmental heterogeneity among distinct subregions is crucial for advancing our understanding of human amygdala development and its relationship to risk for psychiatric disorders (MacMillan et al., 2003; Plessen et al., 2006; Qin et al., 2014; Rosso et al., 2005).

The arborization, plasticity, and migration of late-maturing paralaminar neurons to neighboring nuclei during the pubertal period further indicate that childhood and adolescence are a critical period of change in amygdala substructure (Avino et al., 2018; Page et al., 2022). Recent advancements in MRI amygdala segmentation methods have enabled the study of subregional amygdala development in humans. As such, we recently discovered distinct associations between age and sex for BLN, CEN, and superficial group (Campbell et al., 2021) and associations of CEN volumes with obesity indicators (Kim et al., 2020) in pediatric samples. However, these findings are limited by wide age ranges and small sample sizes. These findings indicate the need for further research to more accurately characterize age-related anatomical changes within structurally and functionally distinct amygdala subregions and how other developmental factors (i.e., sex, pubertal stage, and childhood obesity) influence amygdala composition during preadolescence.

### Rationale

Here, we aimed to characterize amygdala total volumes, subregion volumes, and relative subregion apportionment in a large, diverse population of preadolescents. We leveraged the *CIT168* probabilistic atlas and high-quality 3T MRI data from 3,953 9- and 10-year-olds from the landmark Adolescent Brain Cognitive DevelopmentL (ABCD®) Study. Based on prior literature (Campbell et al., 2021; Dima et al., 2022; Herting et al., 2018; Kim et al., 2020; Page et al., 2022), we hypothesized age- and sex-related differences in amygdala subregions consistent with prior findings and obesity-related differences in CEN volumes. Given prior findings, after accounting for age, we did not expect to find subregion volumes to relate to puberty (Russell et al., 2021).

## Methods and Materials

### Study Population and Dataset

Cross-sectional data from the ABCD® Study were obtained from the baseline visit from the annual 3.0 (MRI data) and 5.0 data release (all other variables) (http://dx.doi.org/10.15154/8873-zj65). The ABCD Study is the largest study of childhood neurodevelopment, enrolling 11,880 children 9 and 10 years of age across 21 sites in the United States between 2016 and 2018 and longitudinally following these adolescents for ten years (Garavan et al., 2018; Volkow et al., 2018). Exclusion criteria for the ABCD Study included lack of English proficiency, severe neurological, medical, or intellectual limitations, and inability to complete an MRI scan (Palmer et al., 2022). The institutional review board (IRB) and human research protection programs at the University of California San Diego oversee all experimental and consent procedures with local IRB approval at the 21 ABCD sites. Each participant provided written assent to participate in the study, and their legal guardian provided written consent. Given that the *CIT168 atlas* was developed and validated on Siemens MRI data (Tyszka and Pauli, 2016) and there are notable between-scanner effects within the ABCD dataset (Dudley et al., 2023), we chose *a priori* to perform amygdala segmentation only on participants collected at 13 of the 21 ABCD sites using Siemens scanners (**Supplemental Methods**). Participants were excluded if their data failed to meet the raw T1w or T2w quality control inclusion standards of the ABCD Consortium (Hagler et al., 2019), had incidental radiological findings (Li et al., 2021), failed *CIT168* segmentation or contrast to noise quality control, or lacked essential covariate data (**Supplemental Figure 1)**. Lastly, to address within-family correlation, we restricted our sample to one randomly selected child per family. We had a final analytic sample of 3,953 (demographic characteristics in **Table 1**).

**Table 1:**
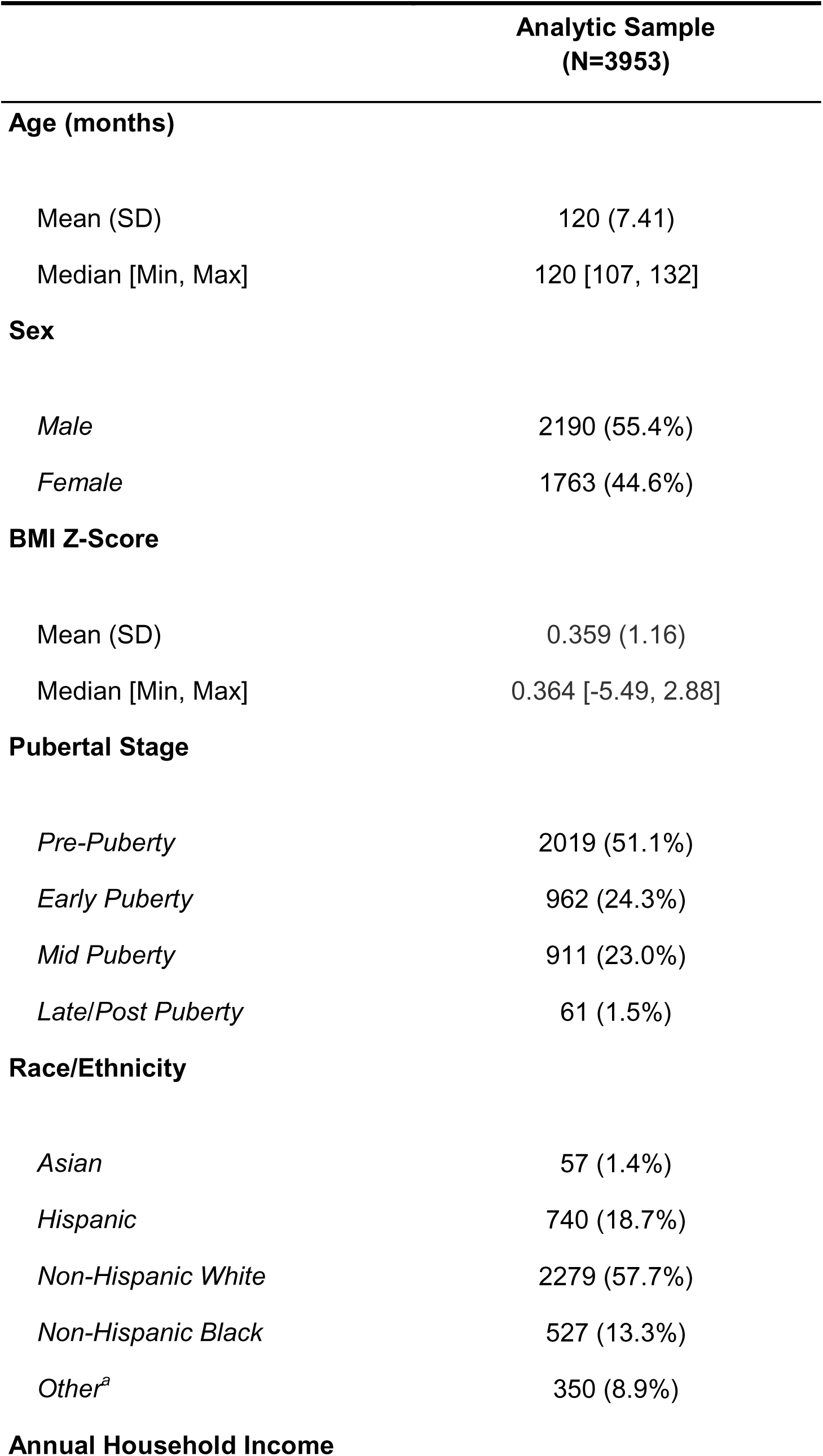
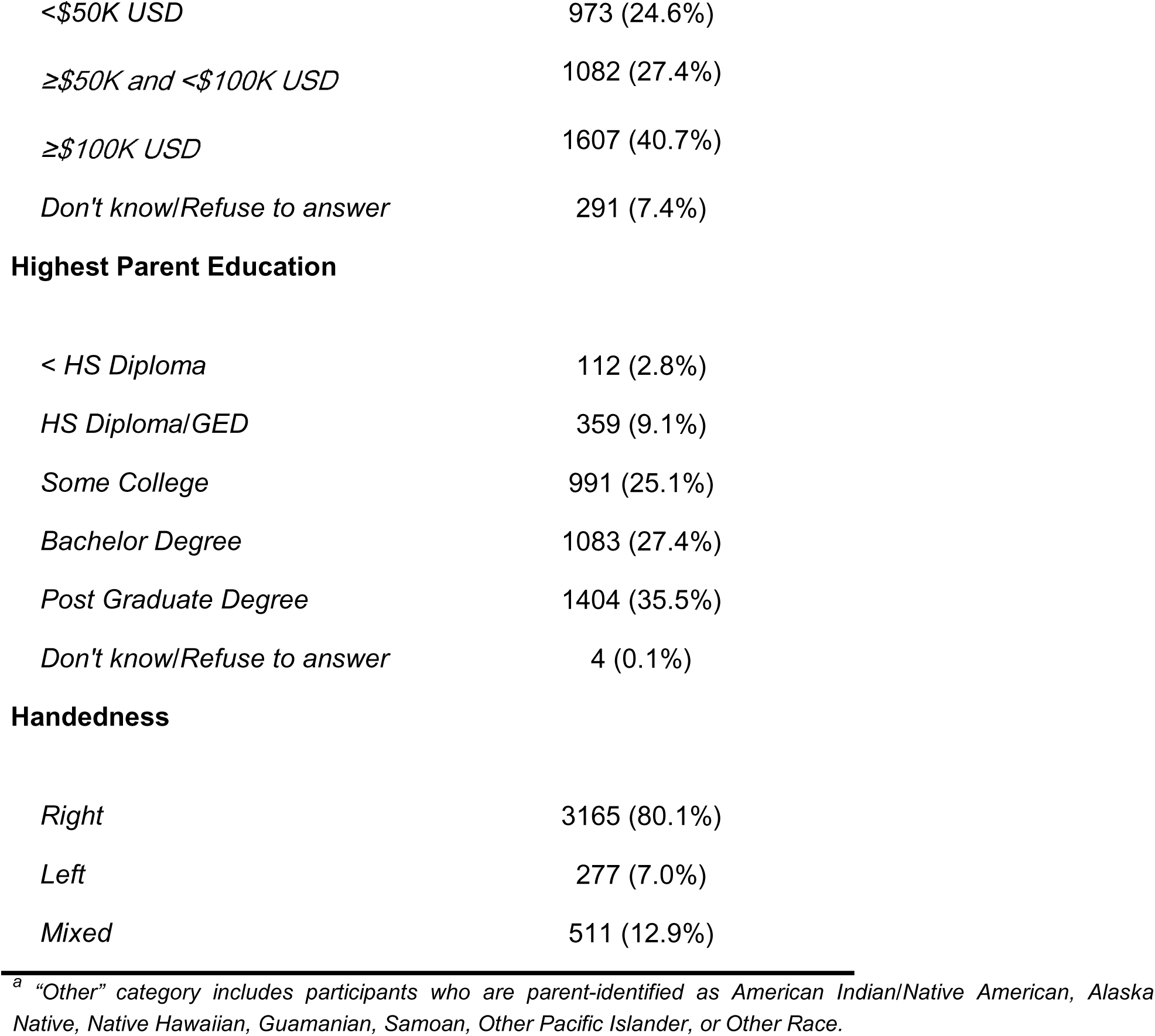
Sample Demographics.

|  | **Analytic Sample**<br>**(N=3953)** |
| --- | --- |
| **Age (months)** |  |
| Mean (SD) | 120 (7.41) |
| Median [Min, Max] | 120 [107, 132] |
| **Sex** |  |
| *Male* | 2190 (55.4%) |
| *Female* | 1763 (44.6%) |
| **BMI Z-Score** |  |
| Mean (SD) | 0.359 (1.16) |
| Median [Min, Max] | 0.364 [-5.49, 2.88] |
| **Pubertal Stage** |  |
| *Pre-Puberty* | 2019 (51.1%) |
| *Early Puberty* | 962 (24.3%) |
| *Mid Puberty* | 911 (23.0%) |
| *Late/Post Puberty* | 61 (1.5%) |
| **Race/Ethnicity** |  |
| *Asian* | 57 (1.4%) |
| *Hispanic* | 740 (18.7%) |
| *Non-Hispanic White* | 2279 (57.7%) |
| *Non-Hispanic Black* | 527 (13.3%) |
| *Other<sup>a</sup>* | 350 (8.9%) |
| **Annual Household Income** |  |
| *<\$50K USD* | 973 (24.6%) |
| *≥\$50K and <\$100K USD* | 1082 (27.4%) |
| *≥\$100K USD* | 1607 (40.7%) |
| *Don't know/Refuse to answer* | 291 (7.4%) |
| **Highest Parent Education** |  |
| *< HS Diploma* | 112 (2.8%) |
| *HS Diploma/GED* | 359 (9.1%) |
| *Some College* | 991 (25.1%) |
| *Bachelor Degree* | 1083 (27.4%) |
| *Post Graduate Degree* | 1404 (35.5%) |
| *Don't know/Refuse to answer* | 4 (0.1%) |
| **Handedness** |  |
| *Right* | 3165 (80.1%) |
| *Left* | 277 (7.0%) |
| *Mixed* | 511 (12.9%) |

<sup>a</sup> "Other" category includes participants who are parent-identified as American Indian/Native American, Alaska Native, Native Hawaiian, Guamanian, Samoan, Other Pacific Islander, or Other Race.

### *CIT168 Amygdala Atlas*: Segmentation and Quality Control

First, unprocessed raw T1- and T2-weighted images collected using harmonized ABCD imaging protocol were downloaded (http://dx.doi.org/10.15154/1520591), and the Human Connectome Project minimal preprocessing pipeline (Glasser et al., 2013) was implemented with registration to the MNI152 template from the FMRIB Software Library (FSL) version 6. Next, the *CIT168 Atlas* (Tyszka and Pauli, 2016) obtained participant-level probabilistic estimates of left and right total amygdala volumes and nine bilateral subregions. The *CIT168 Atlas* groups subregions of the BL based on MRI template visibility. The dorsal and intermediate subdivisions are combined as the BLDI, and the ventral subdivisions are combined with the paralaminar nuclei as the BLVPL. B-spline bivariate symmetric normalization (SyN) diffeomorphic registration algorithm from Advanced Normalization Tools (Avants et al., 2007) version 2.2.0 was adapted for individual image registration to the *CIT168 Atlas* (Figure 1); then, atlas labels were inverse-warped to individual space. Notably, volume estimates using the diffeomorphic registration remain robust even at contrast-to-noise ratios as low as 1.0 (Tyszka and Pauli, 2016). This robustness results from the algorithm incorporating information from a sphere with an approximately 8 mm radius around each voxel when optimizing registration deformations, giving it a significant advantage over the human eye, which relies on 2D sections through the volume. Compared to other atlases, the probabilistic registration approach allows for the preservation of partial volume information and voxel-wise estimations of uncertainty. Using *fslstats* from FSL version 5.0.7, probabilistic volumes for each ROI were calculated by measuring the total number of voxels and multiplying by their mean probability. Based on prior work (Campbell et al., 2022, 2021), only images with all four contrast-to-noise ratios (CNR) ≥1.0 were included in our analyses. Amygdala relative volume fractions (RVFs) were calculated by dividing the volume of a subregion by the total amygdala volume for that hemisphere. Violin plots of subregion volumes are in **Supplemental Figure 2**. To contextualize our estimates of *CIT168 Atlas* amygdala volumes as part of the peer-review process, we also include a comparison with FreeSurfer *Aseg* atlas-derived estimates provided by the ABCD consortium in **Supplemental Materials**.

**Figure 1.**
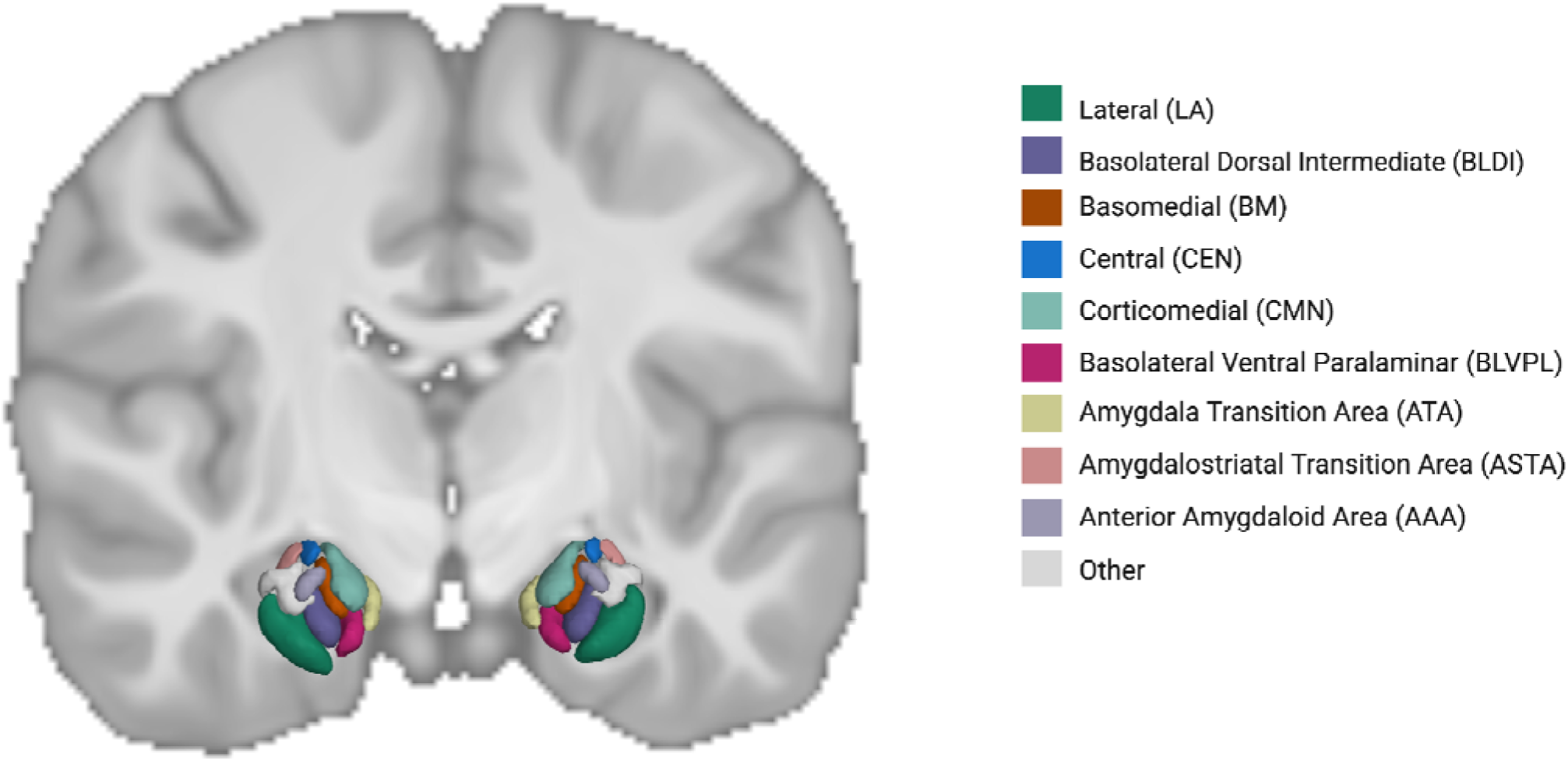
Probabilistic amygdala nuclei segmentation in a representative subject with the *CIT168 Atlas*. Structural T1w and probabilistic estimates of 9 subregions (threshold set at 0.5 for visualization) are shown in the coronal plane.

### Developmental Variables and Covariates

We included the child’s age (*months*), sex at birth (*Male* or *Female*), pubertal stage, and body mass index z-scores (BMIz) as primary predictors of interest based on our hypotheses. We utilized the caregiver report Pubertal Development Scale, which was converted to a corresponding Tanner stage (*Pre-Puberty*, *Early Puberty*, *Mid-Puberty*, *Late Puberty*, and *Post-Puberty*) (Barch et al., 2021; Herting et al., 2020). Due to small categorical numbers, we collapsed the latter two groups (*Late/Post-Puberty*) for analysis. Body mass index (BMI) was calculated in kg/m^2^ using average standing height and weight, measured by a trained research assistant two to three times at the visit. BMIz for sex and age was calculated using the CDC’s United States Growth Charts (Flegal and Cole, 2013). Participants wearing a non-removable prosthetic/cast or having an extreme value (Flegal and Cole, 2013) were excluded.

We included potential confounders associated with amygdala volumes (Brito and Noble, 2014; Dumornay et al., 2023), including race/ethnicity (*Asian, Hispanic, Non-Hispanic White, Non-Hispanic Black, Other racial/ethnic groups* [identified by parents as American Indian/Native American, Alaska Native, Native Hawaiian, Guamanian, Samoan, other Pacific Islander, or other race]), average household income (≥*$100K USD,* ≥*$50k to <$100K USD, <$50K* USD, or *Don’t Know/Refuse to Answer*), and highest household education (*Post-Graduate Degree, Bachelor’s Degree, Some College, High School Diploma/GED, less than High School Diploma,* or *Don’t Know/Refuse to Answer*). Again, due to small categorical numbers, we collapsed the *Asian* and *Other* racial/ethnic groups (*Asian/Other*) for analysis. We included MRI collection precision variables: handedness (*right, left,* or *mixed*) and ABCD site to account for scanner-related differences. In the ABCD dataset, controlling for site effects performs similarly to COMBAT-based harmonization methods in sample sizes larger than 250 subjects (Dudley et al., 2023). We included intracranial volume (ICV) in our volume-based analyses to account for total brain size differences, which are critical in both age- and sex-based analyses during development.

### Statistical Analyses

Analyses were conducted using R Version 4.3.1 (R Core Team, 2023). For Linear Mixed Effect (LME) models, we used *lme4* (Bates et al., 2014). Model diagnostics were assessed using *performance* (Lüdecke et al., 2021). Age was centered at youngest enrollment age, 108 months. To make regional effects directly comparable and aid in the interpretability of parameter estimates, we standardized each MRI outcome (i.e., total amygdala volumes, amygdala subregions volumes, relative proportions of amygdala subregion, ICV) using our analytical sample of 3,953 participants. Standardized parameter estimates are interpreted as the corresponding increase or decrease in standard deviations of the ROI.

Our first LME models measured the effect of our primary developmental variables (i.e., age, sex, pubertal stage, BMIz) on hemispheric amygdala total volume while controlling for covariates (see **Developmental Variables and Covariates**) and random effect of ABCD site. A p-value < 0.05 was used as our threshold of significance. Next, the same LME structure was used with amygdala subregion volumes as the outcome. Lastly, we used LME models to investigate the association with amygdala subregion apportionment using subregion RVF as the outcome. Details in **Supplemental Methods**. Given that RVFs are a proportion of amygdala volume, these models did not include ICV. To account for the multiple comparisons across subregion analyses, false discovery rate (FDR) correction was separately performed for each model set (i.e., volumes, proportions), and an FDR-corrected p-value (p-FDR) < 0.05 was used as our threshold for detecting significant effects. To quantify the overall significance of pubertal stage variable, we used a Type III ANOVA with Satterthwaite’s methods to obtain an F-value and associated p-value.

Thus, volume-based analyses examine the effects of our developmental variables of interest on amygdala subregion volumes relative to brain size. In contrast, RVF-based analyses assess these effects in terms of amygdala subregion volumes as a fraction of amygdala hemispheric volume, addressing questions related to apportionment. Together, this allows us to probe whether a developmental variable is associated with amygdala subregion volume relative to brain size and/or whether it influences the proportional distribution of subregions within the amygdala. This distinction helps clarify whether observed effects reflect global subregion differences, more localized changes in amygdala apportionment, or apparent shifts in amygdala apportionment of smaller subregions that may arise as a reflection of volumetric changes in larger subregions.

## Results

The primary analytic sample was one month older, had more males, a marginally lower BMI z-score, and were more likely to be non-Hispanic white and from higher socioeconomic backgrounds compared to the ABCD Study population (**Supplemental Table 1**).

### Age Effects

Age (in *months*) was positively associated with both total left [β = 0.006, 95% CI: 0.003 – 0.009, p < 0.001] and right amygdala volumes [β = 0.006, 95% CI: 0.003 – 0.009, p < 0.001] (**Supplemental Table 2**). Age was also positively associated with volumes in all amygdala subregions, except for the left AAA and left CMN (**Figure 2** and **Supplemental Table 2**). To demonstrate linearity, we visualized the relationships between age and amygdala volumes using scatterplots fitted with generalized additive models (GAMs) in **Supplemental Figures 3 and 4**. Age was not associated with differences in subregion apportionment except for the left AAA (**Figure 2** and **Supplemental Table 3**).

**Figure 2.**
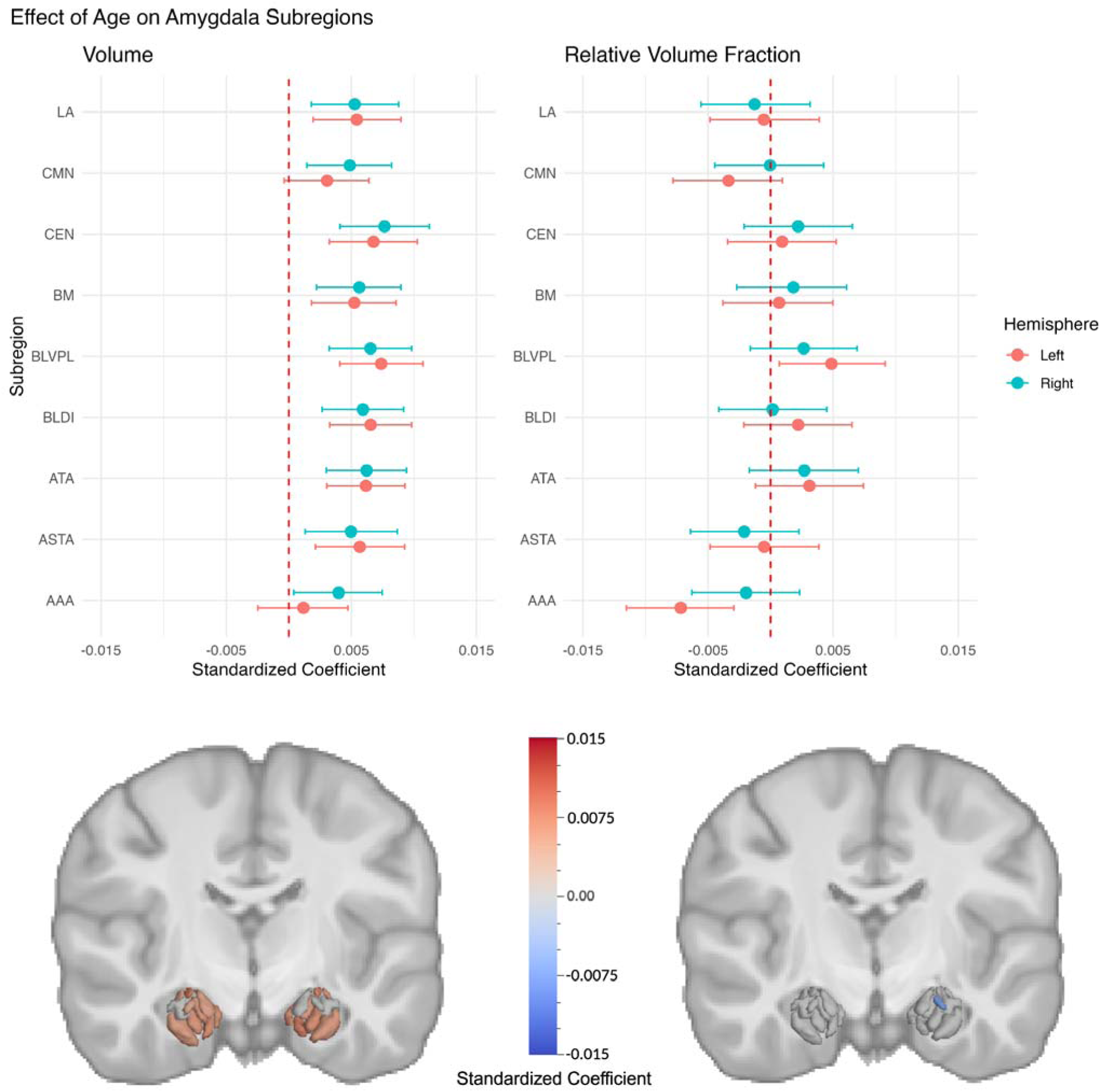
Age Effects on Amygdala Subregions. Standardized beta coefficients with 95% confidence intervals (CIs) for age effects on subregion volumes (left panel) and RVFs (right panel) by hemisphere. CIs intersecting the dashed line indicate null effects. Subregion beta coefficients that remained significant after FDR correction are heat mapped onto the amygdala in the bottom panel of the figure.

### Sex Effects

Female preadolescents had smaller total left [β = -0.233, 95% CI: -0.289 – -0.177, p < 0.001] and right amygdala volumes [β = -0.168, 95% CI: -0.224 – -0.112, p < 0.001] (**Supplemental Table 4**). This sex difference was observed for most amygdala subregions, except for the bilateral CMN, right CEN, and right BM (**Figure 3, Supplemental Table 4**). Sex differences were also observed in the apportionment of amygdala subregions (**Figure 3, Supplemental Table 5**). Bilaterally, females exhibited larger amygdala RVFs in the CMN, CEN, and ASTA, and smaller RVFs in the BLVPL. Females were found to have larger amygdala RVFs in the right BM, but smaller RVFs in the right LA and left ATA.

**Figure 3.**
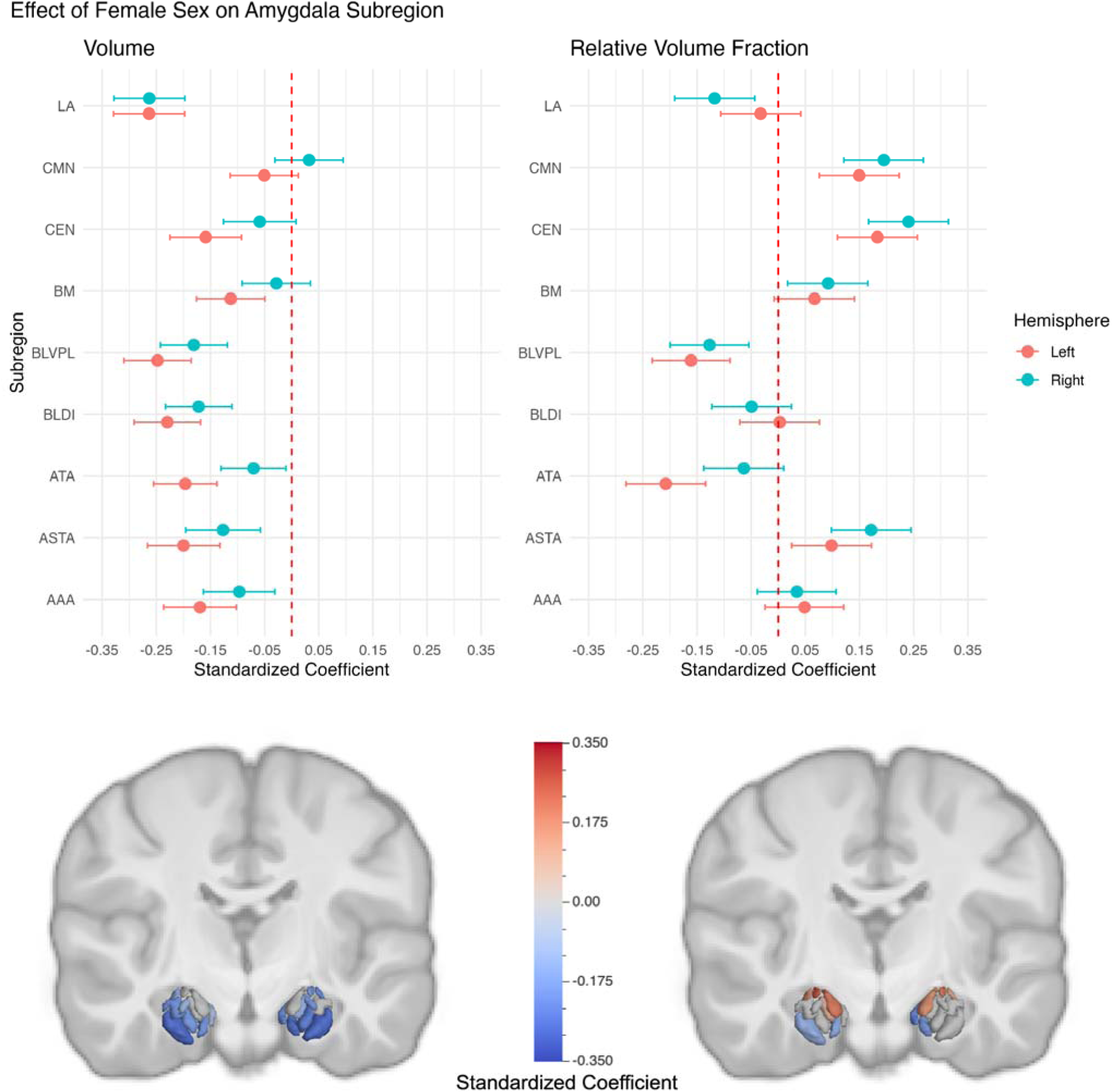
Sex Effects on Amygdala Subregions. Standardized beta coefficients with 95% CIs for female sex effects on subregion volumes (left panel) and RVFs (right panel) by hemisphere. CIs intersecting the dashed line indicate null effects. Subregion beta coefficients that remained significant after FDR correction are visualized on the amygdala in the bottom panel of the figure.

### Pubertal Stage Effects

Pubertal stage was not significantly related to total left [F(3, 3926.5) = 0.246, p = 0.86] or right amygdala volumes [F(3, 3920.5) = 1.876, p = 0.13] (**Supplemental Table 6**). Moreover, pubertal stage was unrelated to subregion volumes or RVFs (**Supplemental Tables 6-7**).

### BMIz Effects

BMIz was negatively associated with total left [β = -0.035, 95% CI: -0.055 – -0.015, p = 0.001] and right amygdala volumes [β = -0.024, 95% CI: -0.044 – -0.004, p = 0.02] (**Supplemental Table 8**). Bilaterally, BMIz was negatively associated with LA and BLDI volumes (**Figure 4, Supplemental Table 8**), and was associated with larger RVFs of the CMN and ATA and smaller RVFs of the LA. Additionally, BMIz was associated with a larger RVF of the right BM and smaller RVF of the right BLDI (**Figure 4, Supplemental Table 9**).

**Figure 4.**
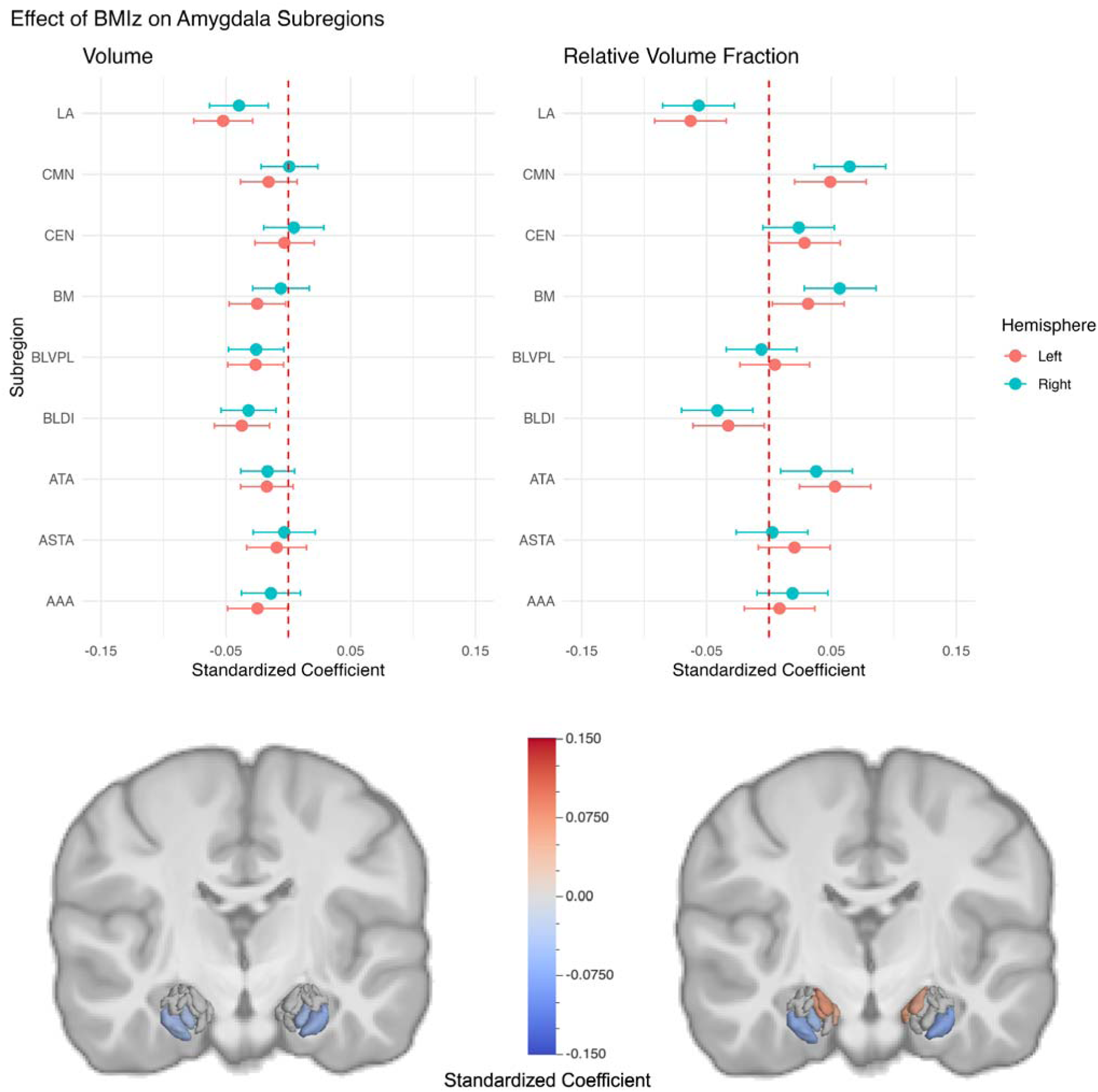
BMIz Effects on Amygdala Subregions. Standardized beta coefficients with 95% CIs for BMIz on subregion volumes (left panel) and RVFs (right panel) by hemisphere. CIs intersecting the dashed line indicate null effects. Subregion beta coefficients that remained significant after correction are visualized on the amygdala in the bottom panel of the figure.

## Discussion

Our findings indicate age, sex, and BMIz are associated with variations in amygdala subregion volumes and apportionment in a large, diverse sample of preadolescents. Age was associated with increases in nearly all subregion volumes, even within this narrow two-year age range; yet age did not explain amygdala apportionment differences. These findings suggest that the amygdala may undergo a near-global (i.e., non-specific) expansion during this period of preadolescence. In RVF analyses, which account for the smaller amygdala volumes in females by measuring subregions as a fraction of hemispheric amygdala volume, sex differences in amygdala apportionment emerged that were consistent across both hemispheres — relatively larger dorsal subregions (i.e., CMN, CEN, ASTA) and relatively smaller BLVPLs in female preadolescents. BMIz, a continuous measure of weight status, was negatively associated with total amygdala volumes, albeit this effect was primarily driven by two large BLN subregions (i.e., LA and BLDI), with relative increases observed in the CMN and ATA in both hemispheres. Lastly, our study failed to detect significant effects of pubertal stage on amygdala volumes or apportionment during this developmental period, contrasting with theorized relationships (Page et al., 2022; Scherf et al., 2013). However, this may be due to (1) an insufficient time window to capture the full effects of puberty, (2) parent-report Tanner Staging method lacking granular resolution, (3) a mismatch between brain and physical pubertal changes, and/or (4) unidentified confounders or moderators. Building upon animal (Ahmed et al., 2008; De Lorme et al., 2012; Malsbury and McKay, 1994), postmortem (Avino et al., 2018), and prior MRI studies (Azad et al., 2021; Campbell et al., 2022, 2021), our findings support the growing evidence that notable differences exist in the structural development of distinct amygdala nuclei and subregions during preadolescence, influenced by age, sex, and indicators of childhood obesity. GWAS studies have revealed that genetic loci associated with volumes of amygdala subregions overlap with genetic risk factors for common brain disorders (Mufford et al., 2023; Ou et al., 2023). Consequently, our findings on amygdala composition provide a foundation for future research investigating how early developmental heterogeneity influences risk and resilience regarding the prevalence and timing of brain disorders that arise during adolescence and adulthood.

Amygdala subregions and their apportionment evolve during development in a sex-specific manner (Campbell et al., 2021). Studying the amygdala as a single, unified structure obscures important individual variability in anatomical changes across its structurally and functionally distinct subregions during childhood and adolescence. The paralaminar nucleus (within the *CIT168* BLVPL subregion) contains neurons that continue to mature and migrate into adulthood (Amaral and Price, 1984; Bernier et al., 2002; deCampo and Fudge, 2012; Tosevski et al., 2002). Notably, many of these quiescent excitatory neurons initiate transcriptional profile changes during adolescence (Sorrells et al., 2019). Some of these neurons migrate to neighboring nuclei, which may contribute to the reconfiguration of amygdala substructure or connectivity (Page et al., 2022), with increases in mature neurons in BLN through middle adulthood (Avino et al., 2018). However, the repository of immature paralaminar neurons in adulthood may serve as a reservoir of amygdalar-hippocampal neuroplasticity (Sorrells et al., 2019). Consistent with other MRI studies (Campbell et al., 2021; Russell et al., 2021), our findings suggest that age, but not puberty, is associated with these changes, with a similar scale of increases across subregion volumes. Age-related increases during 9 and 10 years likely reflect a uniform period of growth before the peak of total amygdala volume. In contrast, prior work observed that non-linear volume decreases within subregions from ages 10 to 17 years occurred in a sex-specific manner (Campbell et al., 2021). Moreover, the amygdala undergoes specific age-related changes in cellular microstructure, with increasing neural density in BLN subregions (i.e., LA, BLDI, BLVPL) (Azad et al., 2021). Future longitudinal study during later developmental periods is warranted to better probe the heterogeneity in amygdala subregional growth patterns following the ‘peak’ period during adolescence and subsequent refinement.

Total amygdala volumes are larger in males, with sex differences in developmental trajectories during adolescence (Dima et al., 2022; Herting et al., 2018). In developing the *CIT168 Atlas*, Tyszka and Pauli (2016) noted sex differences disappeared after normalizing for ICV or amygdala RVF. However, 10- to 17-year-old males had larger absolute volumes of the LA, BLDI, BM, CMN, AAA, and total amygdala when accounting for differences in ICV (Campbell et al., 2021). Here, we confirm the emergence of sex differences as early as ∼10 years of age for most prior identified subregions (e.g., LA, BLDI, AAA, total amygdala) and introduce novel findings (e.g., ASTA, ATA, BLVPL, CEN). Sex differences in amygdalar apportionment patterns observed in the current study were largely consistent (Campbell et al., 2021) — females bilaterally have smaller relative volumes of the BLVPL and larger relative CEN and ASTA. Notably, we observed a novel finding of larger relative CMN for females. These three larger subregions in females are adjacent in the dorsal amygdala, near the stria terminalis — a major efferent pathway of the amygdala that projects to the hypothalamus and modulates the output of its nuclei.(Dudás, 2021) Our novel findings may be attributed to a larger sample size or reflect that sex differences in amygdala subregions vary across development. Interestingly, Campbell et al. (2021) observed age-related changes in male amygdala apportionment led to greater similarity to females by late adolescence.

Previous studies have reported mixed findings between childhood obesity and total amygdala volumes (Jiang et al., 2023; Perlaki et al., 2018; Zaugg et al., 2022). In our study, higher BMIz was associated with smaller total amygdala volumes, primarily driven by smaller BLN subregions (e.g., LA and BLDI), the two largest *CIT168* subregions. After accounting for the smaller amygdala volumes, higher BMIz was linked to decreased relative volumes of these two subregions, and increased relative volumes of the CMN, ATA, and BM. These findings contrast with earlier studies that found no relationship between BMIz and subregions in 405 adolescents (Campbell et al., 2021) and a relationship between CEN volumes and waist-to-height ratio in 71 youth (Kim et al., 2020). While animal models suggest a role of the CEN in regulating homeostatic and cue-mediated feeding (Izadi and Radahmadi, 2022; Petrovich et al., 2009), distinct populations of basolateral principal cells both mediate and suppress appetitive behaviors of the CEN (Kim et al., 2017). Future research is needed to explore whether the observed effects of BMIz relate to higher-order disruption in the eating behavior circuity of the amygdala (Kim et al., 2017). Our work builds upon prior evidence of a relationship between childhood obesity and global amygdala volumes to suggest that pediatric weight status is associated with specific subregional differences. Elucidating the mechanisms linking weight status to amygdala substructure could provide critical insights into the neural underpinnings of disordered eating patterns (e.g., emotional eating behaviors) and inform targeted interventions.

### Future Directions and Limitations

Our study underscores the utility of the *CIT168 Atlas,* a probabilistic atlas of amygdala subregions, in studying neurodevelopment, contributing to a more refined understanding of amygdala substructure development. Importantly, the *in vivo* segmentation template approach uses joint high-accuracy diffeomorphic registration of T1w and T2w structural images to display reliable extraction of major amygdala subregions and has been validated against manual tracing and cross-referencing to four histological sources (Tyszka and Pauli, 2016). While manual tracing is commonly considered the ‘gold standard’ for total amygdala segmentation, its disadvantages include that it is time-consuming and requires operators to have anatomical expertise. Moreover, for large MRI datasets, like the current one, the labor cost of manual tracing is prohibitive and subtle drift in tracing criteria of manual raters while rating large sets of data can occur. Accordingly, we selected to implement *CIT168* over other atlases (such as Freesurfer) as it is a probabilistic segmentation approach, using an atlas developed based on healthy young male and female adults (ages 22-35 years from the human connectome project). The probabilistic segmentation approach, as opposed to deterministic, allows for the preservation of partial volume information and uncertainty estimates (i.e., the potential error is included in each person’s estimate to ‘weight’ their volume rather than treating any given voxel as being part of the amygdala as absolute truth). Additionally, the *CIT168 Atlas* was developed using data from a younger, healthy cohort makes, which reasonably would confer an advantage in studying development compared to atlases, such derived from older and disproportionately male brains (Amunts et al., 2005), such as FreeSurfer. In fact, consistent with prior findings (Morey et al., 2009; Schoemaker et al., 2016; Zhou et al., 2021), the FreeSurfer *Aseg atlas* overestimated left amygdala volumes by 20.8 ± 9.9% (Mean ± SD) and right amygdala volumes by 25.5 ± 9.1% compared to our *CIT168 Atlas* approach in our pediatric sample ages 9- and 10-years-old (**Supplemental Figures 5** and **6**). Importantly, this suggests that the *CIT168 Atlas* may provide a higher accuracy segmentation of total amygdala volumes in pediatric populations. Moreover, the current study used a rigorous approach limited to high-quality 3T images collected from a single manufacturer (Siemens) with appropriate CNRs for high-quality segmentations. However, the final sample did not represent the larger ABCD study cohort, including participants from only 13 of the 21 ABCD study sites. Further research is needed to assess the fidelity of *CIT168 Atlas* registrations to data acquired using other MRI manufacturers. Additionally, our study focuses on a single wave of data from this longitudinal cohort when participants are 9- and 10-years-old. Implementing the atlas in future data waves will be crucial for understanding how age, sex, and body mass influence the developmental trajectories of amygdala subregions throughout adolescence. Lastly, the current study does not examine patterns of amygdala activation using functional MRI or relationships between amygdala subregions and brain disorders in this developmental cohort or, which will be a critical future direction of this work in later developmental waves, as many neuropsychiatric disorders emerge during adolescence (Solmi et al., 2022).

## Conclusions

We uncovered distinct associations between amygdala total volumes, subregion volumes, and subregion relative volumes with age, sex, and BMIz, but not puberty, in nearly 4,000 preadolescents. Age was associated with near-global amygdala growth without changes in apportionment. Female sex was related to smaller total volumes and subregion volumes as well as apportionment differences in dorsal subregions and BLVPL. Our childhood obesity metric was related to smaller total volumes, primarily driven by decreases in two large BLN subregions, which led to greater relative volumes in smaller subregions. This research contributes to the growing evidence that distinct neurodevelopmental patterns exist among heterogenous amygdala subregions, potentially relevant to socioemotional and physical health.

## Acknowledgments

A special thanks to the participants and families of the ABCD Study. The research described in this article was supported by the National Institutes of Health [NIEHS R01ES032295, R01ES031074, NIDA U01DA041048].

Data used in the preparation of this article were obtained from the Adolescent Brain Cognitive Development (ABCD) Study (https://abcdstudy.org), held in the NIMH Data Archive (NDA). This is a multisite, longitudinal study designed to recruit more than 10,000 children aged 9-10 and follow them over 10 years into early adulthood. The ABCD Study is supported by the National Institutes of Health Grants [U01DA041022, U01DA041028, U01DA041048, U01DA041089, U01DA041106, U01DA041117, U01DA041120, U01DA041134, U01DA041148, U01DA041156, U01DA041174, U24DA041123, U24DA041147]. A full list of supporters is available at https://abcdstudy.org/nih-collaborators. A listing of participating sites and a complete listing of the study investigators can be found at https://abcdstudy.org/principal-investigators.html. ABCD consortium investigators designed and implemented the study and/or provided data but did not necessarily participate in the analysis or writing of this report. This manuscript reflects the views of the authors and may not reflect the opinions or views of the NIH or ABCD consortium investigators. The ABCD data repository grows and changes over time.

We would like to acknowledge ABCD Consortium staff for collecting data. We would also like to acknowledge J. Max Landau and Jade Li for their assistance in performing a manual visual quality control of a sample of image segmentations. Alethea de Jesus contributed to data cleaning and formatting of ABCD tabulated data prior to data analyses.

## Author Contributions

LNO, JMT, and MMH conceived and designed the project. ABCD Consortium staff acquired data. LNO, CT, and JM completed the preprocessing and processing of MRI data. LNO, HA, and MMH completed statistical analyses. LNO and MMH drafted the manuscript. All study authors reviewed and edited the manuscript before submission.

