## Supplemental Information for "Amygdala Subregion Volumes and Apportionment in Preadolescents — Associations with Age, Sex, and Body Mass Index"

This document includes:

- Supplemental Methods
- Supplemental Figure 1 Flowchart of subject selection and corresponding N’s
- Supplemental Figure 2 Violin Plot of Amygdala Subregion Volumes
- Supplemental Figure 3 Scatterplots of Left Amygdala Subregion Volumes by Age
- Supplemental Figure 4 Scatterplots of Right Amygdala Subregion Volumes by Age
- Supplemental Figure 5 Scatterplot of Total Left Amygdala Volumes using FreeSurfer *Aseg* and *CIT168* Atlases
- Supplemental Figure 6 Scatterplot of Total Right Amygdala Volumes using FreeSurfer *Aseg* and *CIT168* Atlases
- Supplemental Table 1 Descriptive Statistics of Analytic Sample, Processed CIT168 Sample, and ABCD Cohort
- Supplemental Table 2 Age Effects on Amygdala Subregion Volumes
- Supplemental Table 3 Age Effects on Amygdala Subregion RVFs
- Supplemental Table 4 Female Sex Effects on Amygdala Subregion Volumes
- Supplemental Table 5 Female Sex Effects on Amygdala Subregion RVFs
- Supplemental Table 6 Puberty Effects on Amygdala Subregion Volumes
- Supplemental Table 7 Puberty Effects on Amygdala Subregion RVFs
- Supplemental Table 8 BMIz Effects on Amygdala Subregion Volumes
- Supplemental Table 9 BMIz Effects on Amygdala Subregion RVFs

**Supplemental Methods**

### *ABCD Structural MRI: Acquisition and Quality Control*

A harmonized data protocol was utilized across sites with 3T MRI scanners. Motion compliance training, as well as real-time, prospective motion correction, was used to minimize the impact of head motion on the BOLD signal. T1w images were acquired using a magnetization-prepared rapid acquisition gradient echo (MP-RAGE) sequence, and T2w images were obtained using a fast spin echo sequence with a variable flip angle. Both consist of 176 slices with 1 mm^3^ isotropic resolution. Additional details on scanning protocol are provided by Casey et al. (2018). ABCD study staff visually rated T1w and T2w images for quality control, and we only included data that met minimum quality control standards for recommended inclusion and absence of clinical findings (Hagler 2019) in our image processing pipeline.

*Statistical Model Equations:*

Eq 1. Total Amygdala Volumes:

Std. Right/Left Amygdala Volume ROI ~ sex + age + BMIz + Puberty Stage + Race/Ethnicity + Household Income + Parent Education + Handedness + Std. ICV + (1|ABCD Site)

Eq 2. Amygdala Subregion Volumes:

Std. Subregion Volume ROI ~ Sex + Age + BMIz + Puberty Stage + Race/Ethnicity + Household Income + Parent Education + Handedness + Std. ICV + (1|ABCD Site)

Eq 3. Amygdala Subregion Relative Proportions:

Std. Subregion Relative Proportion ~ Sex + Age + BMIz + Puberty Stage + Race/Ethnicity + Household Income + Parent Education + Handedness + (1|ABCD Site)

**Supplemental Figure 1.** Flowchart of subject selection detailing inclusion and exclusion criteria applied to result in our primary sample.

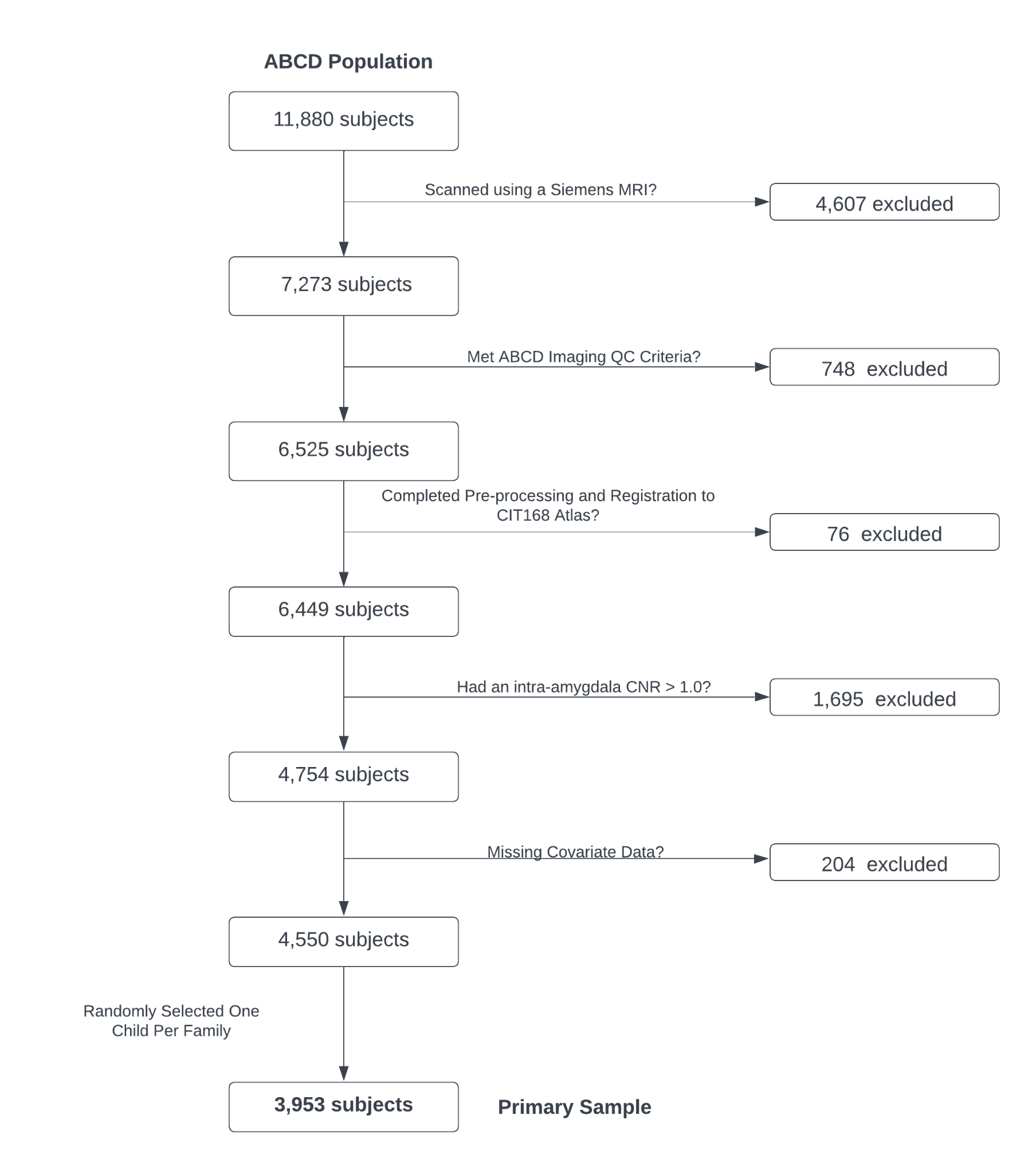

**Supplemental Figure 2. Violin Plot of Amygdala Subregion Volumes (in mm^3^) by Hemisphere**

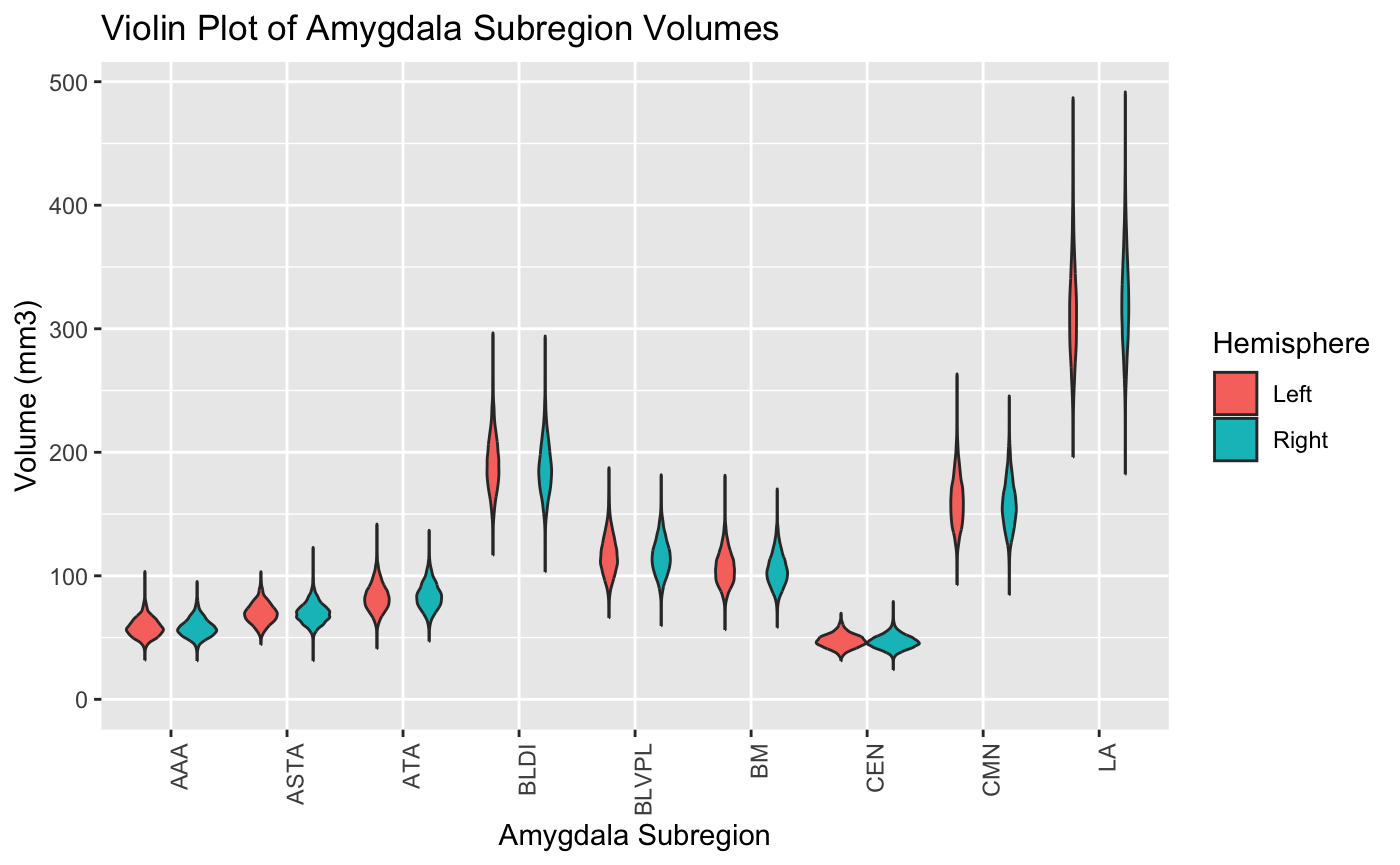

**Figure 3. Scatterplots of Amygdala Subregion Volume (in mm^3^) by Age in the Left Hemisphere.** Trend lines (dark blue) with 95% confidence interval (orange) were fit using generalized additive models (GAMs) to demonstrate grossly linear relationships between age and subregion volumes.

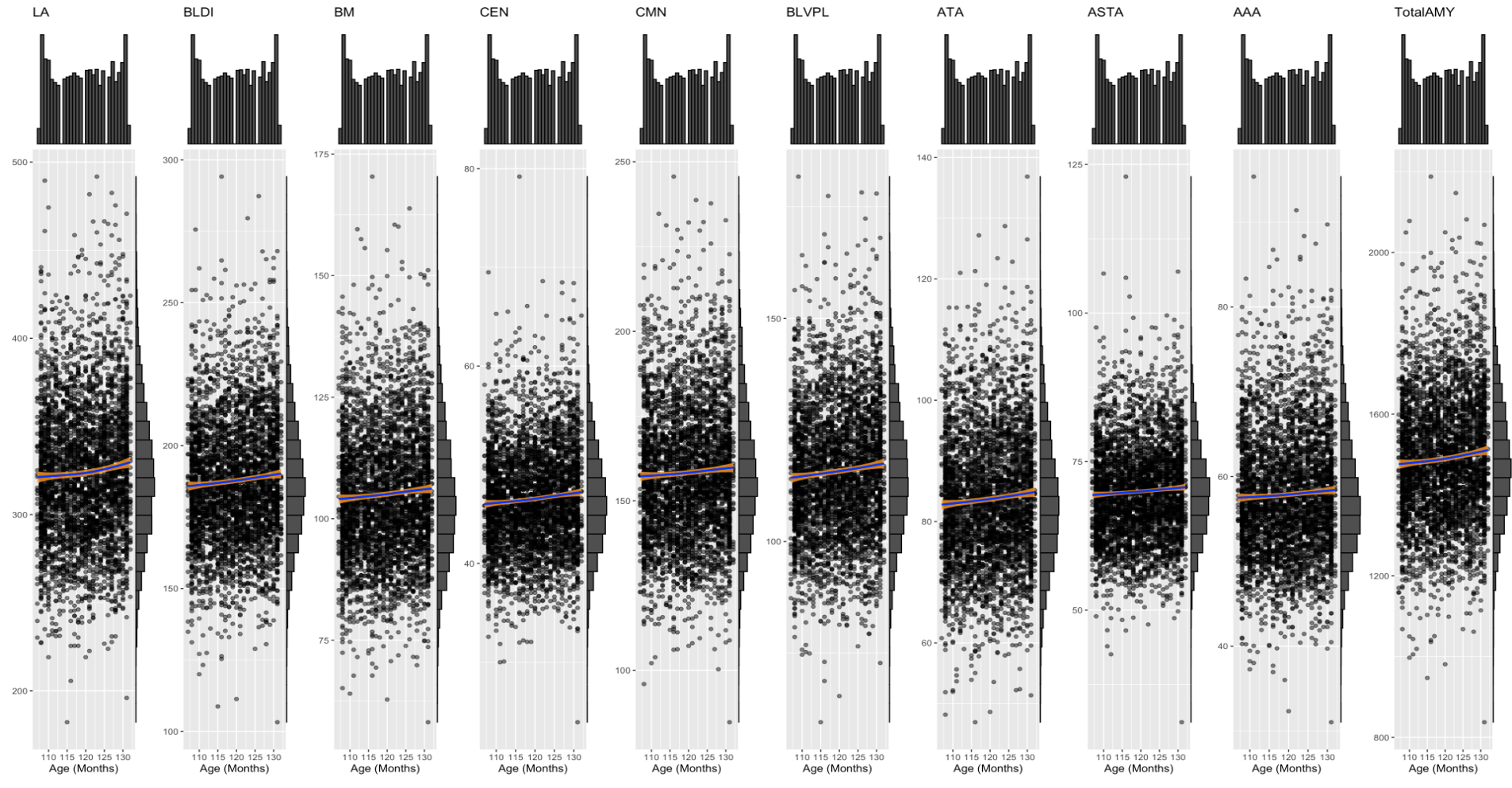

**Figure 4. Scatterplots of Amygdala Subregion Volume (in mm^3^) by Age in the Right Hemisphere.** Trend lines (dark blue) with 95% confidence interval (orange) were fit using generalized additive models (GAMs) to demonstrate grossly linear relationships between age and subregion volumes.

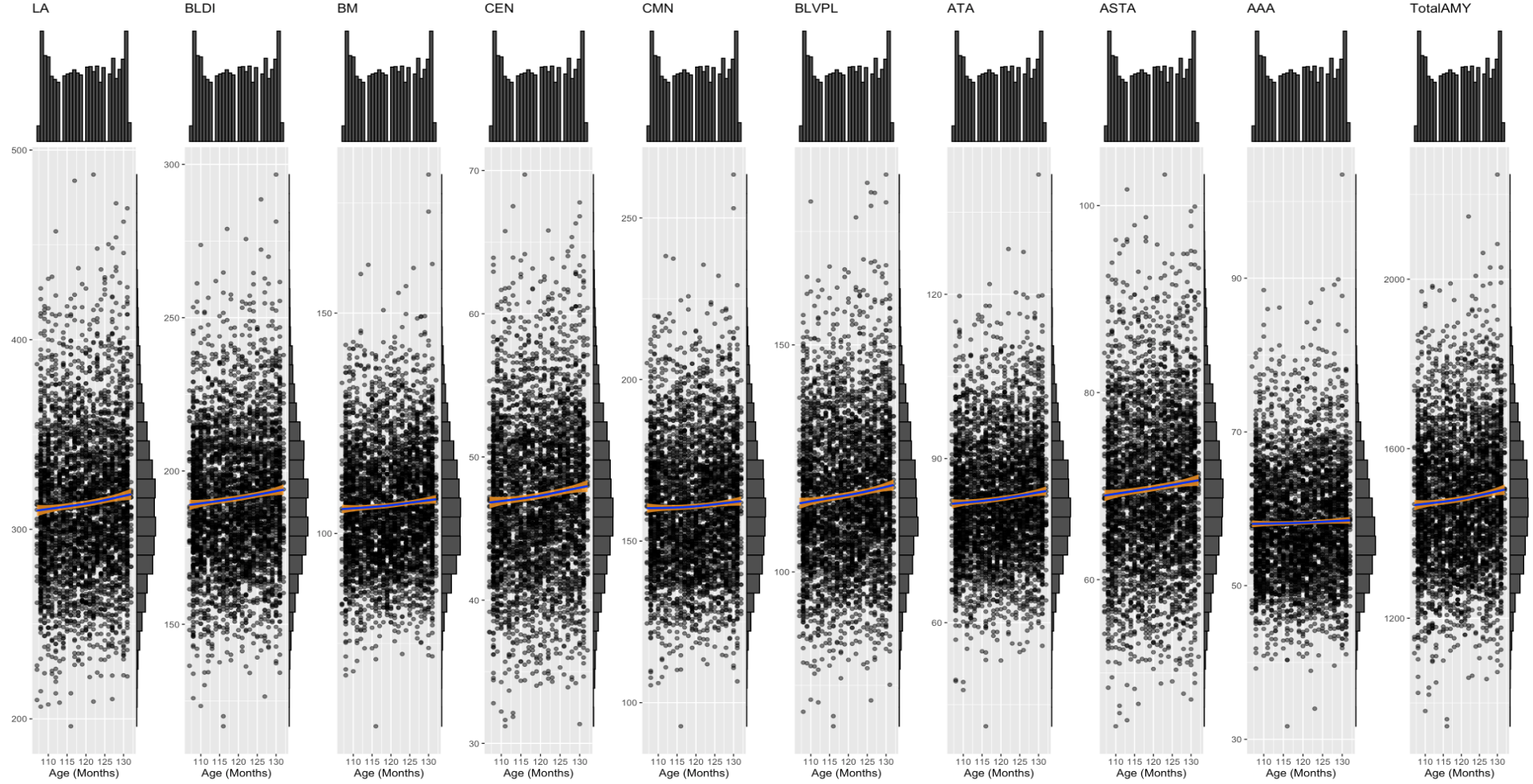

**Figure 5. Comparison of Total Left Amygdala Volume using the *CIT168* Atlas to the FreeSurfer *Aseg* Atlas.** A scatterplot comparing the Total Left Amygdala Volume (mm^3^) using the *CIT168* Atlas and the FreeSurfer *Aseg* Atlas by ABCD site with linear regression lines and confidence intervals demonstrating volume overestimation using FreeSurfer.

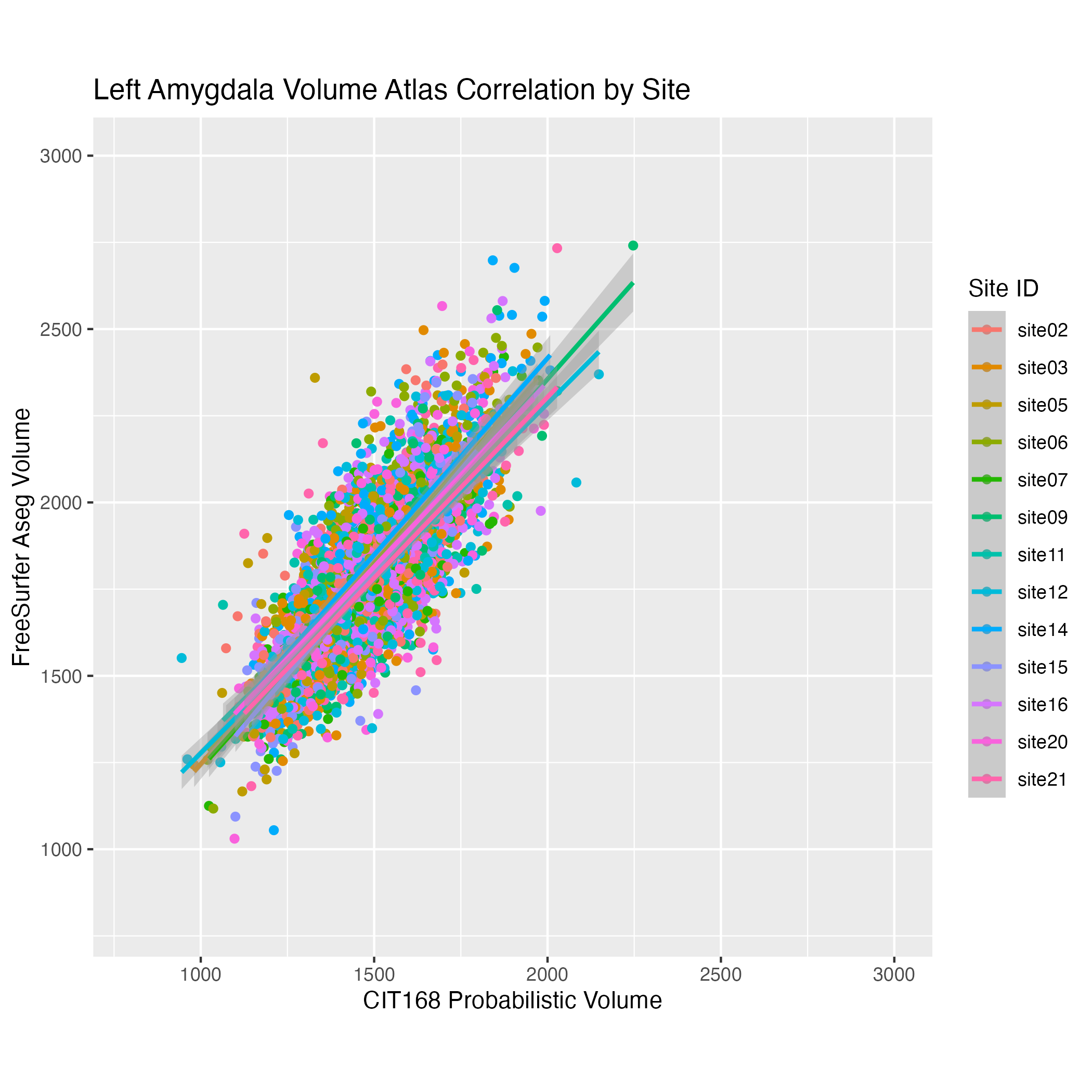

**Figure 6. Comparison of Total Right Amygdala Volume using the *CIT168* Atlas to the FreeSurfer *Aseg* Atlas.** A scatterplot comparing the Total Right Amygdala Volume (mm^3^) using the *CIT168* Atlas and the FreeSurfer *Aseg* Atlas by ABCD site with linear regression lines and confidence intervals demonstrating volume overestimation using FreeSurfer.

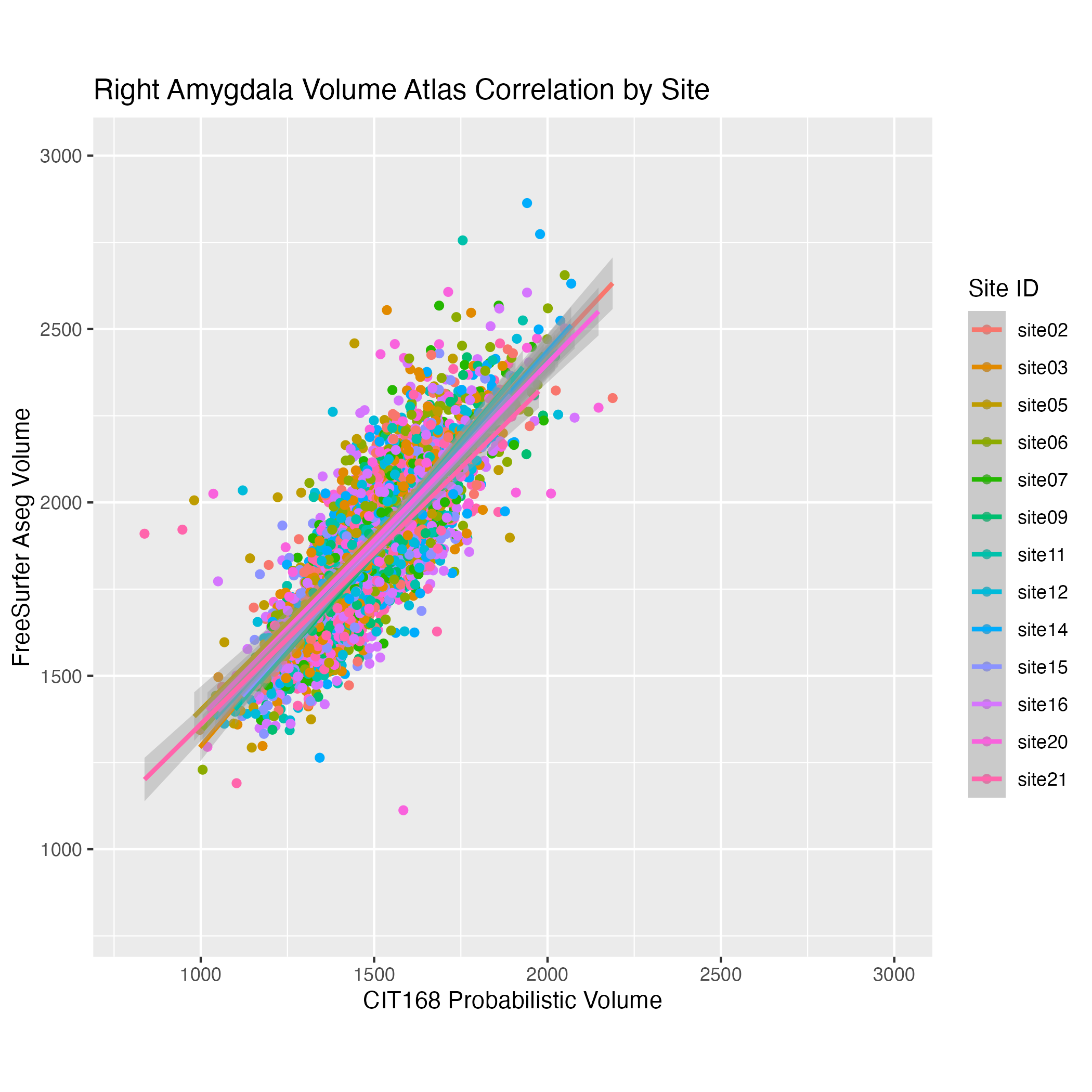

**Supplemental Table 1.** Descriptive Statistics of Primary Sample vs. Processed Sample vs. ABCD Study Population

|  | **Analytic Sample (N=3953)** | **Processed Sample (N=6449)** | **ABCD Cohort**  **(N=11,868)** |
| --- | --- | --- | --- |
| **Age (months)** |  |  |  |
| Mean (SD) | 120 (7.41) | 119 (7.47) | 119 (7.50) |
| Median [Min, Max] | 120 [107, 132] | 119 [107, 132] | 119 [107, 133] |
| Missing | 0 (0%) | 0 (0%) | 1 (0.0%) |
| ***Sex*** |  |  |  |
| Male | 2190 (55.4%) | 3354 (52.0%) | 6188 (52.1%) |
| Female | 1763 (44.6%) | 3094 (48.0%) | 5677 (47.8%) |
| Intersex-Male | 0 (0%) | 1 (0.0%) | 3 (0.0%) |
| **BMIz** |  |  |  |
| Mean (SD) | 0.359 (1.16) | 0.352 (1.18) | 0.407 (1.16) |
| Median [Min, Max] | 0.364 [-5.49, 2.88] | 0.366 [-5.92, 2.88] | 0.413 [-5.92, 3.07] |
| Missing | 0 (0%) | 32 (0.5%) | 77 (0.6%) |
| ***Puberty Stage*** |  |  |  |
| Pre Puberty | 2019 (51.1%) | 3150 (48.8%) | 5837 (49.2%) |
| Early Puberty | 962 (24.3%) | 1475 (22.9%) | 2709 (22.8%) |
| Mid Puberty | 911 (23.0%) | 1487 (23.1%) | 2672 (22.5%) |
| Late/Post Puberty | 61 (1.5%) | 99 (1.5%) | 181 (1.5%) |
| Missing | 0 (0%) | 238 (3.7%) | 469 (4.0%) |
| ***Race/Ethnicity*** |  |  |  |
| White | 2279 (57.7%) | 3562 (55.2%) | 6173 (52.0%) |
| Black | 527 (13.3%) | 1065 (16.5%) | 1784 (15.0%) |
| Hispanic | 740 (18.7%) | 1146 (17.8%) | 2410 (20.3%) |
| Asian | 57 (1.4%) | 83 (1.3%) | 252 (2.1%) |
| Other | 350 (8.9%) | 592 (9.2%) | 1247 (10.5%) |
| Missing | 0 (0%) | 1 (0.0%) | 2 (0.0%) |
| ***Household Income*** |  |  |  |
| <$50K USD | 973 (24.6%) | 1646 (25.5%) | 3222 (27.1%) |
| ≥$50K and <$100K USD | 1082 (27.4%) | 1762 (27.3%) | 3068 (25.9%) |
| ≥$100K USD | 1607 (40.7%) | 2550 (39.5%) | 4561 (38.4%) |
| Don't know/Refuse to answer | 291 (7.4%) | 491 (7.6%) | 1015 (8.6%) |
| Missing | 0 (0%) | 0 (0%) | 2 (0.0%) |
| ***Parent Education*** |  |  |  |
| < HS Diploma | 112 (2.8%) | 239 (3.7%) | 593 (5.0%) |
| HS Diploma/GED | 359 (9.1%) | 574 (8.9%) | 1132 (9.5%) |
| Some College | 991 (25.1%) | 1706 (26.5%) | 3074 (25.9%) |
| Bachelor Degree | 1083 (27.4%) | 1739 (27.0%) | 3013 (25.4%) |
| Post Graduate Degree | 1404 (35.5%) | 2182 (33.8%) | 4042 (34.1%) |
| Missing/Refused | 4 (0.1%) | 9 (0.1%) | 14 (0.1%) |
| ***Handedness*** |  |  |  |
| Right | 3165 (80.1%) | 5147 (79.8%) | 9372 (79.0%) |
| Left | 277 (7.0%) | 456 (7.1%) | 847 (7.1%) |
| Mixed | 511 (12.9%) | 846 (13.1%) | 1590 (13.4%) |
| Missing | 0 (0%) | 0 (0%) | 59 (0.5%) |
| **ICV** |  |  |  |
| Mean (SD) | 1540000 (133000) | 1530000 (136000) | 1490000 (144000) |
| Median [Min, Max] | 1540000 [1070000, 2080000] | 1520000 [1000000, 2080000] | 1490000 [984000, 2080000] |
| Missing | 0 (0%) | 20 (0.3%) | 140 (1.2%) |

**Supplemental Table 2. Age Effects on Total Amygdala and Subregion Volumes**

Standardized beta-coefficient, confidence intervals, P-value, and FDR-corrected P-Values for the effect of age (in months) on total amygdala and subregion volumes in the left and right hemispheres.

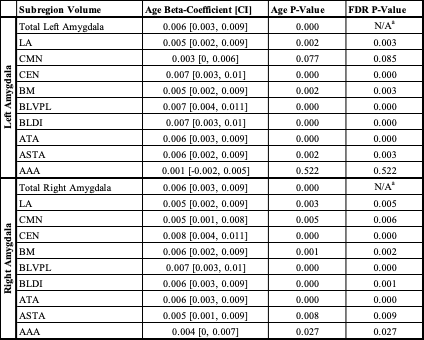

^a^ N/A = Not Applicable; FDR correction was not performed for Eq 1. Total Amygdala Volumes.

**Supplemental Table 3. Age Effects on Amygdala Subregion RVFs**

Standardized beta-coefficient, confidence intervals, P-value, and FDR-corrected P-Values for the effect of age (in months) on subregion RVFs in the left and right amygdala.

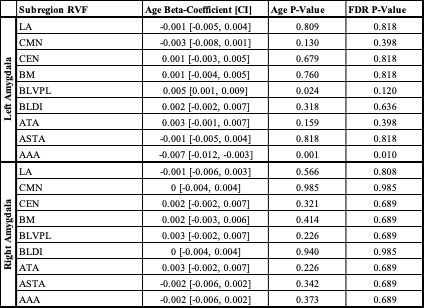

**Supplemental Table 4. Sex Effects on Total Amygdala and Subregion Volumes**

Standardized beta-coefficient, confidence intervals, P-value, and FDR-corrected P-Values for the effect of female sex on total amygdala and subregion volumes in the left and right hemispheres.

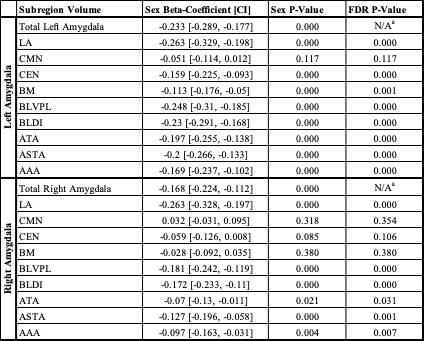

^a^ N/A = Not Applicable; FDR correction was not performed for Eq 1. Total Amygdala Volumes.

**Supplemental Table 5. Sex Effects on Amygdala Subregion RVFs**

Standardized beta-coefficient, confidence intervals, P-value, and FDR-corrected P-Values for the effect of female sex on subregion RVFs in the left and right amygdala.

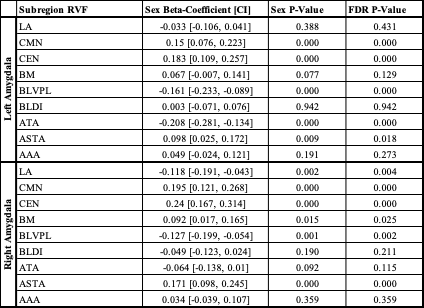

**Supplemental Table 6. Puberty Status Effects on Total Amygdala and Subregion Volumes**

Type-III ANOVA F-Value, P-value, and FDR-corrected P-Values for the cumulative effect of pubertal status on total amygdala and subregion volumes in the left and right hemispheres.

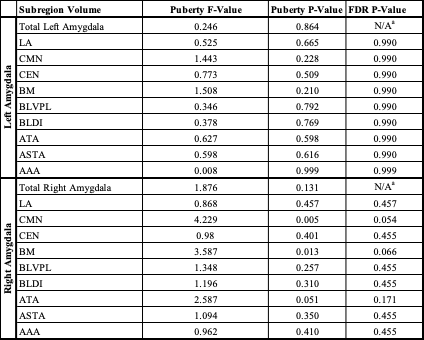

^a^ N/A = Not Applicable; FDR correction was not performed for Eq 1. Total Amygdala Volumes.

**Supplemental Table 8. Puberty Status Effects on Amygdala Subregion Volumes.**

Type-III ANOVA F-Value, P-value, and FDR-corrected P-Values for the cumulative effect of puberal status on subregion RVFs in the left and right amygdala.

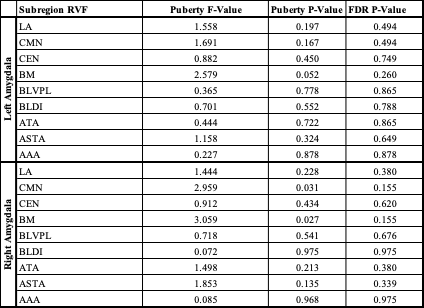

**Supplemental Table 8. BMIz Effects on Total Amygdala and Subregion Volumes.**

Standardized beta-coefficient, confidence intervals, P-value, and FDR-corrected P-Values for the effect of BMIz on total amygdala and subregion volumes in the left and right hemispheres.

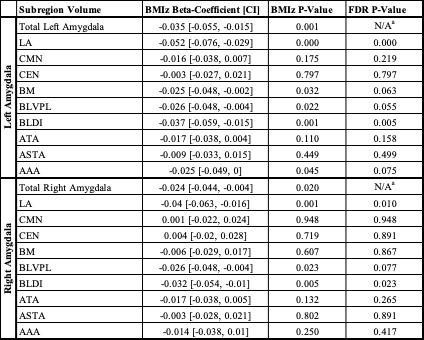

### ^a^ N/A = Not Applicable; FDR correction was not performed for Eq 1. Total Amygdala Volumes.

### **Supplemental Table 9. BMIz Effects on Amygdala Subregion RVFs**

Standardized beta-coefficient, confidence intervals, P-value, and FDR-corrected P-Values for the effect of BMIz on subregion RVFs in the left and right amygdala.

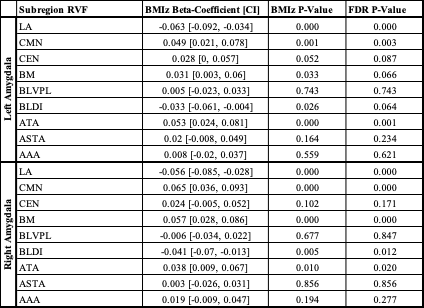
